# Do the loops in the N-SH2 binding cleft truly serve as allosteric switch in SHP2 activation?

**DOI:** 10.1101/2020.11.18.388447

**Authors:** Massimiliano Anselmi, Jochen S. Hub

## Abstract

SHP2 is a critical regulator of signal transduction implicated in developmental disorders and cancer. SHP2 is activated by phosphopeptide binding to the N-SH2 domain, triggering the release of N-SH2 from the catalytic PTP domain. Based on early crystallographic data, it has been widely accepted that opening of the binding cleft of N-SH2 serves as a key “allosteric switch” for SHP2 activation. We critically review structural data and use extensive molecular simulations to test the “allosteric switch” model of activation. We find that the binding cleft in N-SH2 is constitutively flexible and open in solution, and that a closed cleft found in certain structures has been imposed by crystal contacts. The free energy cost for N-SH2 release is only marginally influenced by the binding cleft. We conclude that not the N-SH2 binding cleft but instead the opening of a central *β*-sheet of N-SH2 is the key allosteric switch triggering SHP2 activation.

## INTRODUCTION

Src-homology-2-containing protein tyrosine phosphatase 2 (SHP2), encoded by the *PTPN11* gene, is a classical non-receptor protein tyrosine phosphatase (PTP). It has emerged as a key downstream regulator of several receptor tyrosine kinases (RTKs) and cytokine receptors, functioning as positive or negative modulator in multiple signaling pathways [1–4]. Germline mutations in the human *PTPN11* gene have been associated with Noonan syndrome and with Noonan syndrome with multiple lentigines (formerly known as LEOPARD syndrome), two multisystem developmental diseases [5–16]. Somatic *PTPN11* mutations were also linked with several types of human malignancies [17–25], such as myeloid leukemia [7, 26–30].

The structure of SHP2 includes two tandemly-arranged Src homology 2 domains (SH2), called N-SH2 and C-SH2, followed by the catalytic protein tyrosine phosphatase (PTP) domain, and a C-terminal tail with a poorly characterized function (Figure 1) [34]. The SH2 domains are structurally conserved recognition elements [35] that allow SHP2 to bind signaling peptides containing a phosphorylated tyrosine (pY) [36]. The N-SH2 domain consists of a central antiparallel *β*-sheet, composed of three *β*-strands, denoted *β*B, *β*C and *β*D, flanked by two *α*-helices, denoted *α*A and *α*B (Figure 1A). The peptide binds in an extended conformation to the cleft that is perpendicular to the plane of the *β*-sheet [31]. N-SH2 contains a conserved affinity pocket covered by the BC loop (also called phosphate-binding loop or pY loop), whose interaction with pY increases the binding of the peptide by 1000-fold relative to unphosphorylated counterparts [37]. Residues downstream to the pY bind to a more variable, less conserved site, which confers binding specificity and which is flanked by the EF and BG loops [38].

**Figure 1:**
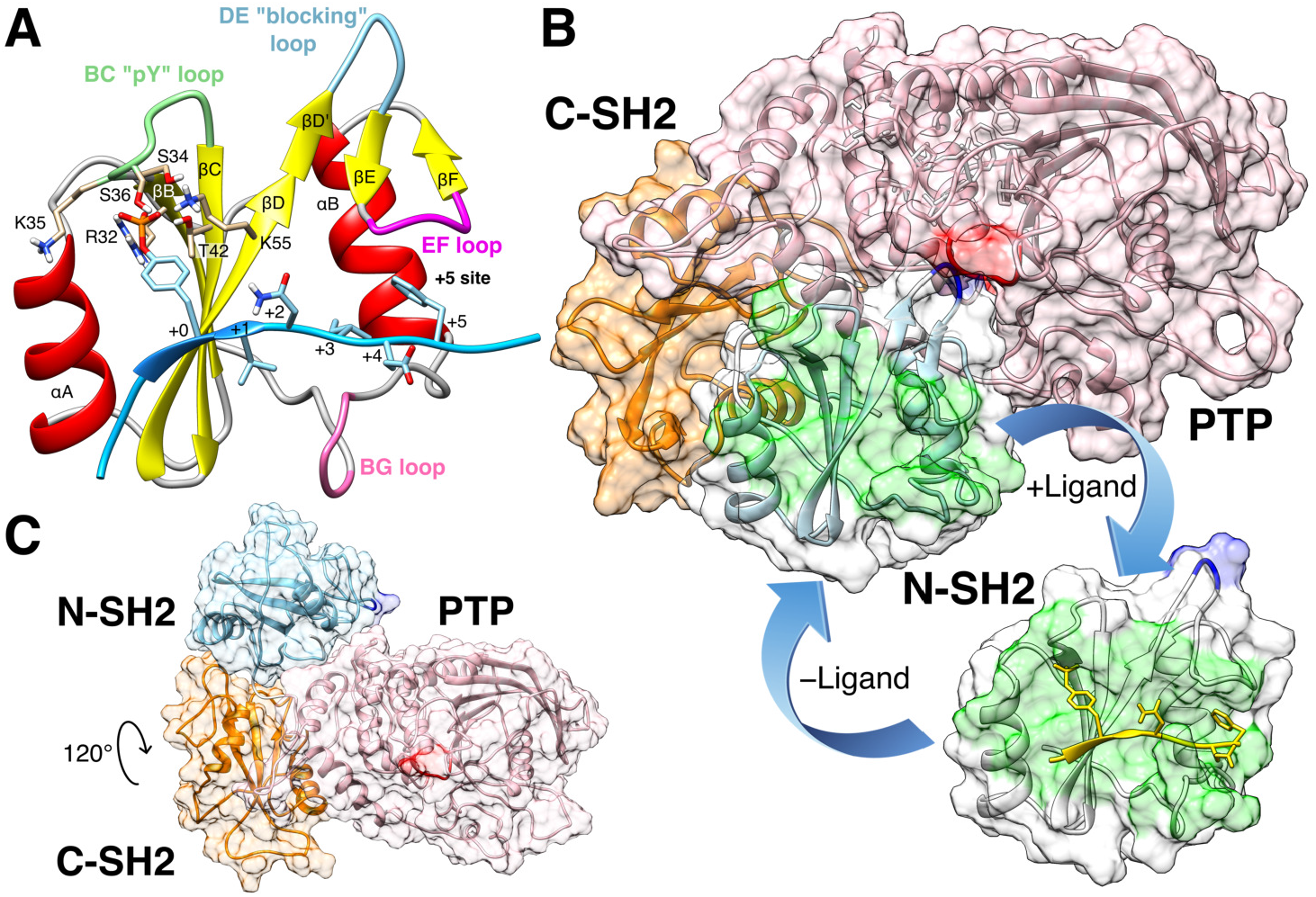
(A) Cartoon representation of the N-SH2 domain in complex with the IRS-1 pY895 peptide [31]. The peptide, shown in cyan, comprises the phosphotyrosine and residues from position +1 to +5 relative to the phosphotyrosine (see labels). Functionally important loops are highlighted in color: BC “pY” loop (green), DE “blocking loop” (light blue), EF loop (magenta), BG loop (deep pink). The phosphotyrosine binds the site delimited by the pY loop and the central *β*-sheet (*β*B, *β*C, βD strands). EF and BG loops delimit the binding cleft (+5 site), where the peptide residue in position +5 is settled. (B) Crystal structure of autoinhibited SHP2 [32]: the N-SH2 domain (cyan cartoon) blocks the catalytic site (red) of the PTP domain (pink) with the blocking loop (blue). The N-SH2 domain is connected to PTP in tandem with the homologous C-SH2 domain (orange). Closure of the N-SH2 binding cleft (green region), delimited by the EF and BG loops, precludes high-affinity phosphopeptide binding. According the “allosteric switch” model, the change in shape of the N-SH2 domain, that accompanies binding of phosphopeptide (yellow), perturbs surface complementarity for the PTP active site, thus promoting N-SH2 dissociation from the PTP domain. (C) Crystal structure of SHP2^E76K^ [33]: the open conformation reveals a 120° rotation of the C-SH2 domain, relocation of the N-SH2 domain to a PTP surface opposite the active site, and a solvent-exposed catalytic pocket.

In 1998, the first crystallographic structure of SHP2 at 2 Å resolution (pdbID 2SHP) revealed that the N-SH2 domain tightly interacts with the PTP domain (Figure 1B), so that the DE loop of N-SH2, thereafter indicated as “blocking loop”, occludes the active site of PTP, forcing SHP2 into an autoinhibited “closed” state [32]. SHP2 structures of the active “open” state, obtained for the basally active, leukemia-associated E76K mutant, showed an alternative relative arrangement of N-SH2 and PTP, that exposed the active site of the PTP domain to the solvent (Figure 1C) [33]. Unexpectedly, in both the autoinhibited and the active state of SHP2, the N-SH2 ligand binding site is exposed to solvent and does not directly interact with PTP or other domains [32, 33, 39]. Hence, the need for an allosteric mechanism was proposed, according to which the binding of a phosphopeptide triggers a series of structural rearrangements in the N-SH2 domain to drive its release from PTP and the consequent activation of SHP2 [32, 39].

The comparison of the autoinhibited structure of SHP2 (pdbID 2SHP) [32] with the existing structures of the isolated N-SH2 domain, either in the absence of (pdbID 1AYD) [31] or in complex with a phosphopeptide (pdbID 1AYA, 1AYB, 4QSY) [31], showed that the EF and BG loops of the N-SH2 domain may undergo large conformational changes (Figure 1B). In the autoinhibited structure of SHP2, the N-SH2 domain shares surface complementarity with the PTP catalytic site, but, as a result of the displacement of the EF loop towards the BG loop, it also contains a closed binding cleft that renders the N-SH2 domain unable to accommodate the C-terminal part of the phosphopeptide, in contrast to the isolated N-SH2 domain, that exhibits an open binding cleft. Therefore, peptide binding seems only compatible with the conformation of isolated N-SH2, but not with the conformation of N-SH2 in autoinhibited SHP2 [32, 39].

Because (i) the closure of the binding cleft in N-SH2 has been ascribed to its interaction with PTP, and (ii) the conformation adopted by the EF loop correlates with the activation of SHP2 in available structural data, the EF loop has been suggested as the key “allosteric switch” that drives the release of N-SH2 from PTP [32, 39]. In light of that, conformation selection [9, 21,40] and induced fit [41] models were put forward for the molecular events leading to functional activation of SHP2; however, both models consider the conformational change involving the EF loop as the key mechanism that drives SHP2 opening [9, 21,40,41]. In conclusion, it has been widely accepted that the N-SH2 binding cleft, delimited by the EF and the BG loop, plays the role of an allosteric switch for the activation of SHP2 [39, 42]. Accordingly, the binding of a ligand at the binding cleft of the N-SH2 domain would induce a transmitted conformational change that prevents PTP domain binding on the other side of N-SH2, and *vice versa* [32, 39].

However, the “allosteric switch” model does not explain how the signal, coming from the displacement of the EF loop, is propagated to the rest of the protein [39]. In addition, considering that the EF loop might be flexible, the comparison of crystallographic structures does not provide the energetic penalties involved in the motion of the EF loop and, consequently, the degree of destabilization of the N-SH2/PTP complex upon the binding cleft opening [39]. Hence, the role of the EF loop as the key allosteric switch has not been rationalized in energetic terms.

Recently, an allosteric interaction in N-SH2 has been proposed as an alternative mechanism of SHP2 activation [43]. Molecular dynamics (MD) simulations have shown that the N-SH2 domain may adopt two distinct conformations, denoted as *α*- and *β*-state, which differ primarily in the conformation of the central *β*-sheet. In the *α*-state, the central *β*-sheet is open, adopting a Y-shaped structure; in the *β*-state, the central *β*-sheet is closed, adopting a parallel structure. The MD simulations suggested that the *β*-state of N-SH2 stabilizes the N-SH2/PTP contacts and, hence, the autoinhibited SHP2 conformation. In contrast, the *α*-state drives the N-SH2 dissociation and SHP2 activation. Notably, the *α*–*β* model of activation rationalized modified basal activity and responsiveness to ligand stimulation of certain mutations at codon 42 [15]. However, the *α*–*β* model seems to contrast the previously suggested “allosteric switch” model.

To resolve this discrepancy, we revisited the “allosteric switch” model. We critically reviewed available crystallographic data of SHP2 in the autoinhibited state. In addition, we used MD simulations, free energy calculations, and enhanced sampling techniques to reveal the conformational dynamics of the binding cleft delimited by the BG and the EF loop in the isolated N-SH2 domain in water, in SHP2 in water, and in SHP2 in a crystal environment. Our results suggest that the binding cleft of N-SH2 is constitutively flexible and the effect of its degree of opening on the activation of SHP2 is negligible. In addition, free energy calculations revealed that, in the crystal environment, the closure of the binding cleft is not due to the allosteric interaction with the PTP domain, but instead a result of the crystal contacts affecting the binding cleft conformation.

## RESULTS

### N-SH2 domain loops are characterized by structural disorder

In crystal structures, the EF and the BG loop have been generally modeled such that the binding cleft of the N-SH2 domain is inaccessible to the ligand [16, 32]. To test whether the closed binding cleft is supported by available data, we first evaluated Ramachandran outliers, electron densities, residue-wise crystallographic R-factors (Z-scores), as well as crystal contacts in the loop regions. The evaluation of the first resolved crystallographic structure of autoinhibited SHP2 (2SHP) [32], performed by MolProbity [44], revealed that the BG loop (residues 84-98) adopts a configuration compatible with a closed binding cleft only at the cost of significant energetic penalties. In particular, the Ramachandran angles of the residues Asn^92^ (*ϕ* = 58.4°; *ψ* = 118.9°) and Asp^94^ (*ϕ* = −42.0°; *ψ* = 95.5°) represent outliers, being represented in less than 0.01% of the samples in protein databank; in case of His^84^ (*ϕ* = −145.7°; *ψ* = 64.1°), Glu^90^ (*ϕ* = −87.7°; *ψ* = −109.2°), and Gly^93^ (*ϕ* = 73.9°; *ψ* = −46.0°), the conformations are allowed yet disfavored, being represented in only 1.24 %, 0.07 % and 1.4 % of the samples, respectively. In the light of the “allosteric switch” model, these unfavorable conformations may be justified considering that the N-SH2 domain structure is forced by PTP in a tensed conformation, which may be turned into a relaxed conformation only upon ligand binding or upon the release from PTP [32,39–41]. However, inspection of the electron density map showed that the electron densities at BG loop, EF loop and pY loop were much lower compared to the density of the nearby *β*-strands (Figure S1), indicating increased disorder, and therefore a reduced structural definition of these loops.

The 2SHP model was refined to a crystallographic R-factor of 0.2 with a R_free_ of 0.27 [32], characterizing a structure with overall acceptable quality [45]. However, the R-factor is a global measure of the model accuracy. In proteins, some regions and strands may be intrinsically disordered such that they cannot be resolved by x-ray diffraction irrespective on the resolution and the overall quality of the experimental data. For this reason, the Z-score has been used as a measure of local model accuracy [46, 47]. The Z-score is given by *Z* = (*R* — 〈*R*〉_res_)/*σ*_res_ [46–48], where 〈*R*〉_res_ and *σ*_res_ are respectively the expected R-factor and the standard deviation of the R-factor values, calculated with the same amino acid in structures within the same resolution range. For instance, for the case of 2SHP with a resolution of 1.95 Å, the R-factor of each residue is compared with the expected R-factor and standard deviation detected from the same kind of residue in all structures of the database with resolution between 1.8 and 2.0 Å [48]. A large, positive spike in a Z-score plot implies that a residue has an R-factor value that is considerably worse than that of the average residue of the same type in structures determined at similar resolution [48]. Z-scores are considered as outliers if greater than 2 [46, 48].

The structures of the N-SH2 domain belonging to the two SHP2 chains, A and B, in the 2SHP asymmetric unit are depicted in Figure 2A [32]. The backbone is colored according to the Z-score values reported in Figure 2B. Evidently, the regions colored in red corresponding to BG loop and to pY loop are poorly resolved, with Z-score values far above the threshold. Similarly, for the EF loop the Z-scores are barely fair in case of chain B, whereas in chain A the corresponding Z-scores are good.

**Figure 2:**
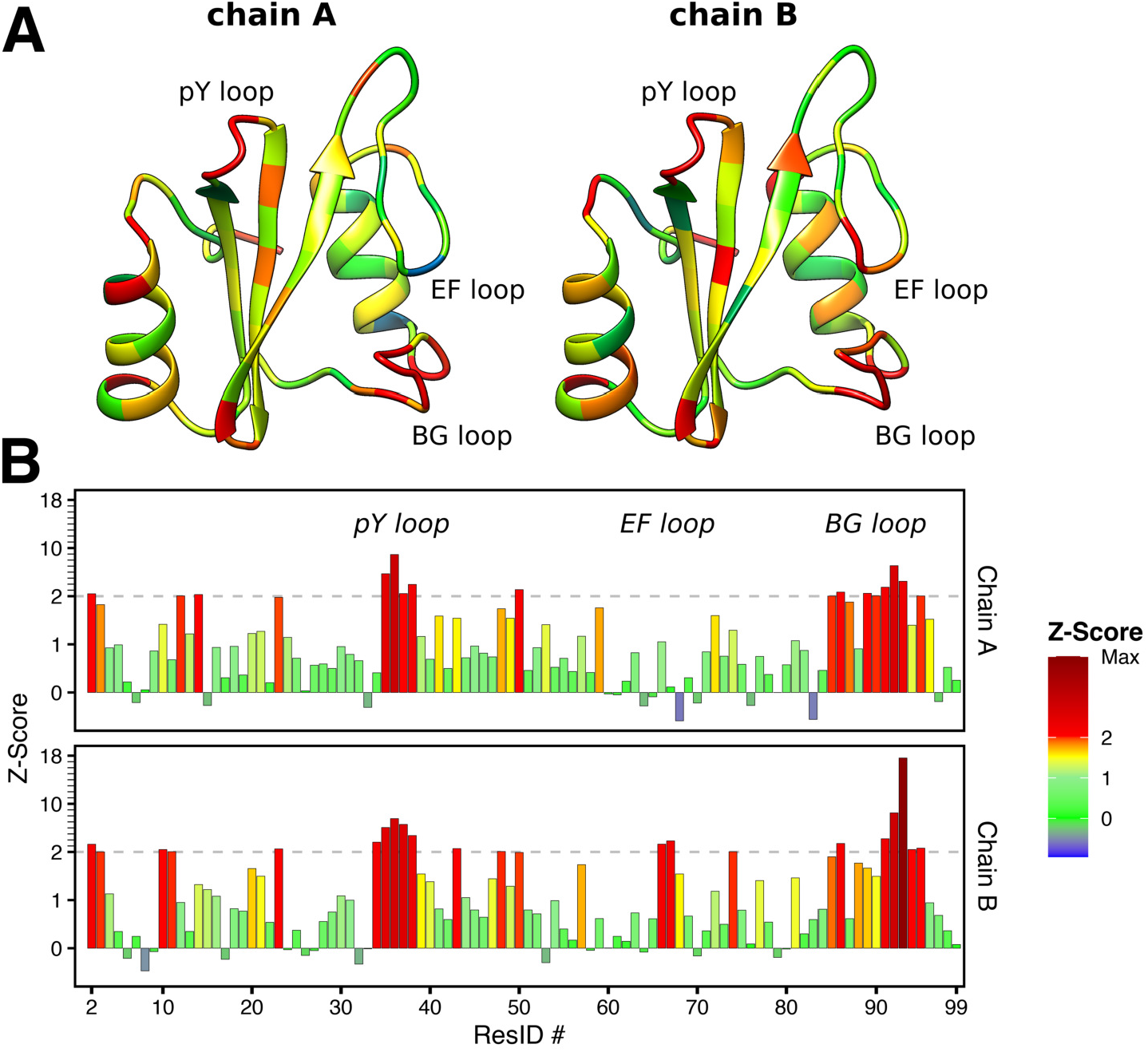
(A) Structures of the N-SH2 domain belonging respectively to the chain A and B of the asymmetric unit reported in the crystal structure of autoinhibited SHP2 (pdbID 2SHP [32]). The backbone is colored on the base of the R-factor Z-scores reported in (B). The outlier threshold of 2 is indicated by dashed lines.

The “allosteric switch” model formulation was based on the conformation adopted by the BG and the EF loop in the 2SHP crystal structure [32], thus the validation of the model would require that those loops be univocally defined. However, the analysis of the electron density map, and the Z-scores reveal that the loops are poorly resolved compared to the rest of the protein. Instead, the loops have been modeled without adequate electron density, and forced in a geometry with significant structural outliers.

### The BG loop is generally poorly resolved in crystal structures of autoinhibited SHP2

Together with the 2SHP structure [32], we computed Z-scores for other, more recent crystal structures of autoinhibited SHP2, comprising either wild type or functional mutants [16,49,50]. Table 1 lists the crystal structures considered in this study, spanning resolutions from 1.87 to 2.7 Å. Figure 3 presents the Z-scores in color scale, for functionally important loop regions, corresponding to the pY loop (Ser^34^–Asp^40^, dark green), the EF loop (Tyr^66^–Glu^69^, purple) and the BG loop (Gly^86^–Glu^97^, pink). In most of the structures, the residues of the BG loop are either missing (blank spaces) or modeled without adequate electron density (yellow-red squares), confirming that BG loop is often disordered and therefore poorly resolved by x-ray diffraction. Likewise, the pY loop is sometimes missing or poorly resolved, although it is less affected by structural disorder than BG loop. In contrast, the short EF loop is marginally affected by structural disorder, being resolved in nearly all crystal structures, although Z-scores for residue Tyr^66^ are often poor.

**Figure 3:**
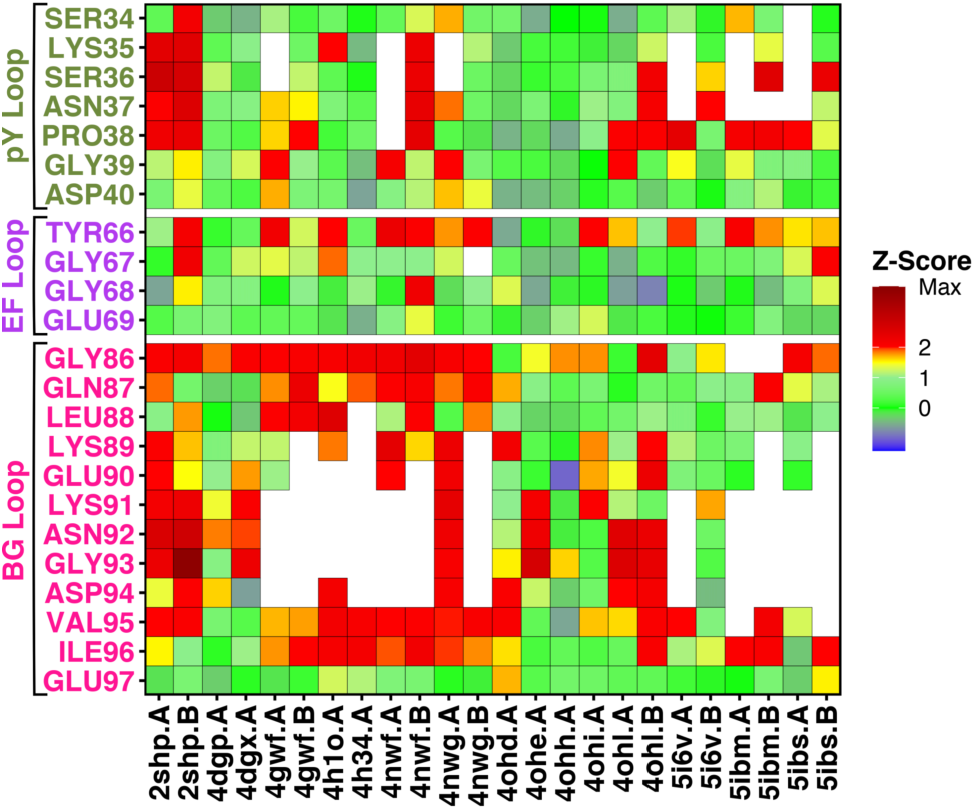
Tile plot of the R-factor Z-scores, calculated for the crystal structures of autoinhibited SHP2. Z-scores are reported in color scale for each residue (blank spaces indicate missing residues), belonging to functionally important loop regions, corresponding to the pY loop (Ser^34^–Asp^40^, dark green), the EF loop (Tyr^66^–Glu^69^, purple) and the BG loop (Gly^86^–Glu^97^, pink).

**Table 1.** Experimental data of autoinibited SHP2 crystal structures.

| PDB id | Space group | Resolution ($\text{\AA}$) | R-factor | R<sub>free</sub> | Description |
| --- | --- | --- | --- | --- | --- |
| 2SHP | P12<sub>1</sub>1 | 2.00 | 0.199 | 0.270 | Wild type |
| 4DGP | P2<sub>1</sub>2<sub>1</sub>2 | 2.30 | 0.199 | 0.255 | Wild type |
| 4DGX | P2<sub>1</sub>2<sub>1</sub>2 | 2.30 | 0.212 | 0.266 | Y279C |
| 4GWF | P12<sub>1</sub>1 | 2.10 | 0.211 | 0.243 | Y279C |
| 4H1O | P2<sub>1</sub>2<sub>1</sub>2 | 2.20 | 0.211 | 0.233 | D61G |
| 4H34 | P2<sub>1</sub>2<sub>1</sub>2 | 2.70 | 0.227 | 0.242 | Q506P |
| 4NWF | P2<sub>1</sub>2<sub>1</sub>2<sub>1</sub> | 2.10 | 0.267 | 0.309 | N308D |
| 4NWG | P2<sub>1</sub>2<sub>1</sub>2<sub>1</sub> | 2.45 | 0.281 | 0.326 | E139D |
| 4OHD | P2<sub>1</sub>2<sub>1</sub>2 | 2.70 | 0.206 | 0.271 | A461T |
| 4OHE | P2<sub>1</sub>2<sub>1</sub>2 | 2.51 | 0.190 | 0.256 | G464A |
| 4OHH | P2<sub>1</sub>2<sub>1</sub>2 | 2.70 | 0.206 | 0.266 | Q506P |
| 4OHI | P2<sub>1</sub>2<sub>1</sub>2 | 2.20 | 0.210 | 0.262 | Q510E |
| 4OHL | P12<sub>1</sub>1 | 2.40 | 0.194 | 0.259 | T468M |
| 5I6V | P12<sub>1</sub>1 | 1.87 | 0.203 | 0.232 | F285S |
| 5IBM | P12<sub>1</sub>1 | 2.18 | 0.192 | 0.240 | S502P |
| 5IBS | P12<sub>1</sub>1 | 2.32 | 0.189 | 0.238 | E76Q |

Taken together, the Z-scores detected in many crystal structures confirm that loops represent flexible regions of the N-SH2 domain, in particular the BG loop. Notable exceptions are a number of high-quality crystal structures (4DGP, 4OHH, 5I6V) [16, 49, 50], in which the loops have been clearly resolved. These structures with well-resolved loops enabled us to further analyze a putative effect of crystal packing on loop conformations.

### N-SH2 interacts with other copies of SHP2 in crystal structures

In the crystal environment, proteins do not adopt the same biological assembly according their physiological conditions, but instead every protein is surrounded by other copies of the same protein, establishing close and rather extensive interactions that stabilize the crystal in a particular space group symmetry. If functionally relevant or even flexible regions are involved in such crystal contacts, the conformations of those regions must be interpreted with care.

Figure 4A shows the surrounding of the N-SH2 domain for the structure 4DGP, which produced good Z-scores even for the BG, EF, and pY loops [16]. Evidently, the N-SH2 domain does not bind only the PTP domain belonging to the same chain (colored in white), but it also interacts with a number of other protein replicas (colored in blue, purple, green and pink). Specifically, the pY, EF, and BG loops, which would be solvent-exposed under physiological conditions, are in contact with other chains. Hence, the conformations of these loops are likely influenced by such interactions, and not exclusively by the binding of the PTP domain on the other face of N-SH2.

**Figure 4:**
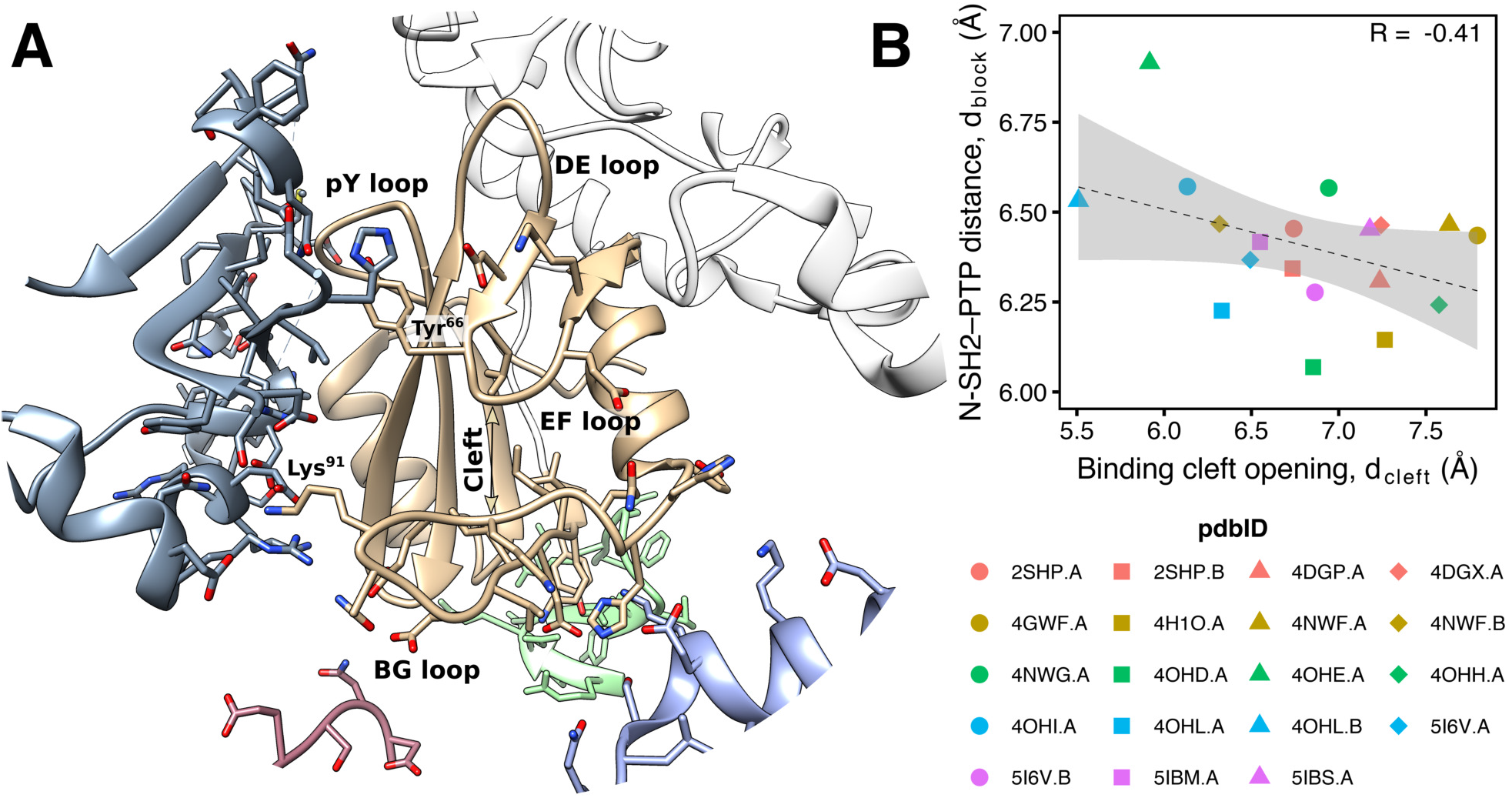
(A) Cartoon representation of the surrounding of the N-SH2 domain in the crystal structure of autoinhibited SHP2 (pdbID 4DGP [16]). PTP domain belonging to the same protein chain is colored in white. Other protein replicas, interacting with the central N-SH2 domain (colored in tan), are depicted respectively in blue, purple, green and pink. (B) Correlation between the N-SH2 binding cleft opening, d_cleft_ (Gly^67^ C_α_–Lys^89^ C_α_ distance) and the N-SH2 blocking loop distance from the catalytic PTP loop, d_block_ (Asp^61^ C_α_–Ala^460^ C_α_ distance), as taken from crystal structures of autoinhibited SHP2, comprising either wild type or functional mutants [16, 32, 49, 50].

In particular, Lys^91^ side chain in BG loop forms salt bridges with charged residues of another replica, whereas the motion of the Tyr^66^ side chain in EF loop is sterically hindered by the presence of other bulky side chains. Even the pY loop, which in absence of a phosphate group is typically modelled in a partially closed conformation, corresponding to an intermediate conformation between the *α*- and *β*-state [43], interacts with acidic side chains of Glu^313^ and Glu^348^ belonging to another protein chain. Therefore, the partially closed loops in the N-SH2 domain might have been stabilized by crystal contacts.

### Crystal packing in autoinhibited SHP2 hinders the N-SH2 motions

According to the “allosteric switch” model, opening of the binding cleft would drive the release of N-SH2 from the PTP domain. Hence, a positive correlation in crystal structures might be expected between (i) cleft opening, given by the Gly^67^ C_*α*_–Lys^89^ C*_α_* distance *d*_cleft_, and (ii) release of the DE blocking loop from the catalytic PTP loop, given by the Asp^61^ C*_α_*–Ala^460^ C*_α_* distance *d*_block_. On the other hand, according to the *α-β* model, a positive correlation might be expected between the opening of the central *β*-sheet and the distance *d*_block_. However, despite a significant spread of *d*_block_ in available crystal structures of autoinhibited SHP2, no such correlation exists. For instance, *d*_cleft_ and *d*_block_ are even (poorly) anticorrelated (Figure 4B), in disagreement with the expectation from the “allosteric switch” model. Also, the spread of the N-SH2 central *β*-sheet lacks the correlation with *d*_block_ (Figure S2).

These data further indicate that crystal packing largely hinders conformational motions of N-SH2 in autoinhibited SHP2 involved in SHP2 activation, and correlations cannot be detected irrespective on the activation mechanism proposed. Hence, to (i) fully resolve the effect of crystal packing on N-SH2 dynamics, and (ii) to quantify the influence of N-SH2 dynamics on SHP2 activation, we employed MD simulations in solution and in a crystal.

### The N-SH2 domain binding cleft is constitutively flexible

According to the “allosteric switch” model, the opening and the closure of the N-SH2 binding cleft is coupled with the activation or the stabilization of autoinhibited SHP2 [32, 39]. Considering that the activation of SHP2 presumably involves a free energy difference of several tens of kJ/mol [41, 43], it is mandatory to evaluate the energy required to open the binding cleft in different conditions.

In order to quantify the intrinsic flexibility of the binding cleft, we used umbrella sampling to compute the potential of mean force (PMF, also referred to as ‘free energy profile’) of the binding cleft opening for the isolated N-SH2 domain (Figure 5A/B). As reaction coordinate, we used the distance *d*_cleft_ defined as the C*_α_*–C_*α*_ distance between Gly^67^ (belonging to the EG loop) and Lys^89^ (belonging to the BG loop). The PMF covers conformations of a completely inaccessible cleft (*d*_cleft_ = 5 Å, Figure 5A cyan cartoon) up to a wide open cleft (*d*_cleft_ = 14 Å, Figure 5A purple cartoon). In Figure 5B, the PMF shows a single free energy minimum at ~11 Å corresponding to an open cleft, in line with the 1AYD crystal structure of apo N-SH2 [31]. This finding is further supported by previous MD simulations of isolated N-SH2 [43], thereby clearly confirming that the cleft is open in the isolated N-SH2 domain.

**Figure 5:**
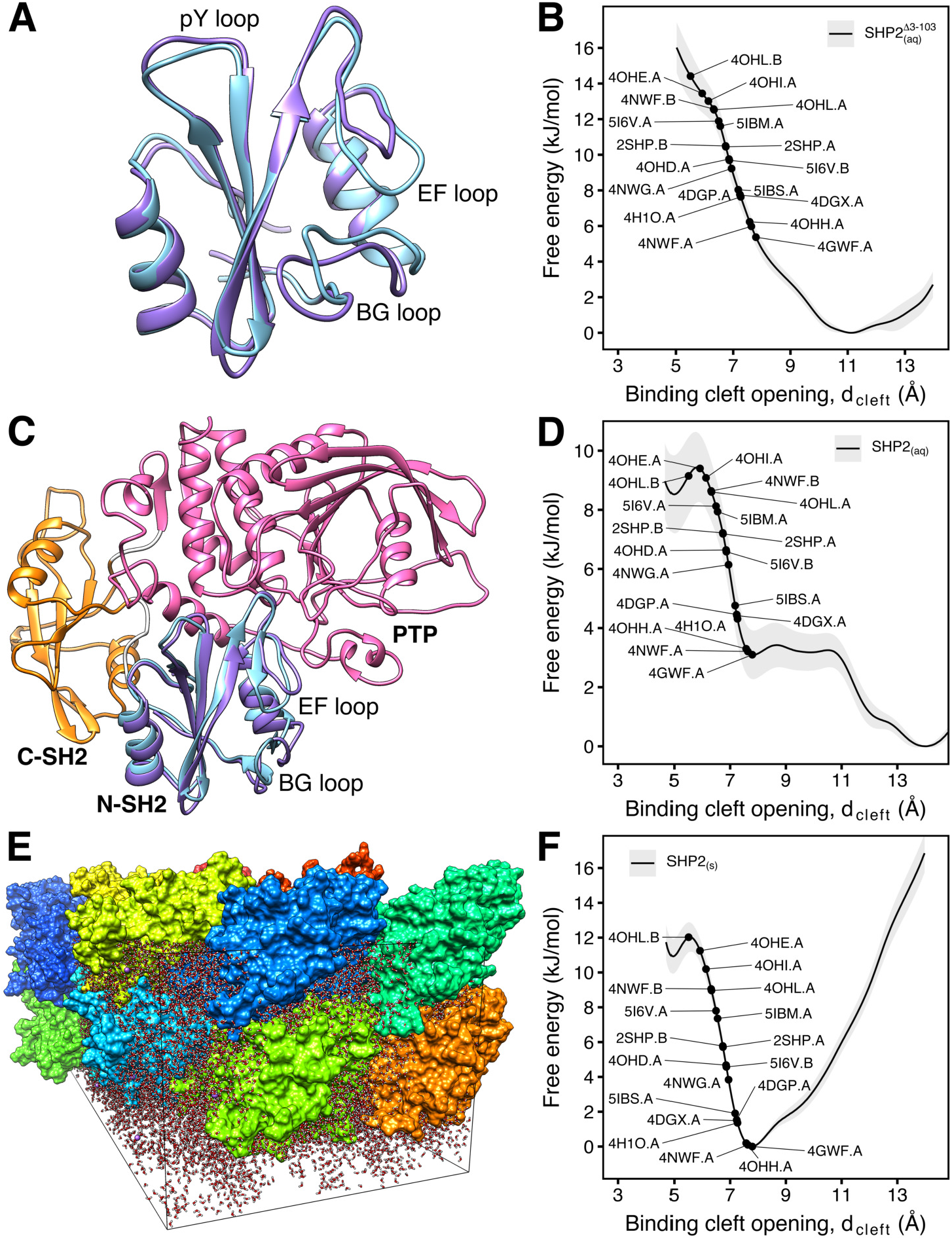
(A) Cartoon representations of the N-SH2 domain in water with the binding cleft either closed (cyan cartoon) or open (purple cartoon). (B) Potential of mean force (PMF) of the binding cleft opening, d_cleft_, for the N-SH2 domain in water. Black dots indicate the degree of cleft opening found in various crystal structures. (C) Cartoon representations of autoinhibited SHP2 in water with the binding cleft either closed (cyan cartoon) or open (purple cartoon). (D) PMF of the binding cleft opening, d_cleft_, for autoinhibited SHP2 in water. (E) Crystal structure of autoinhibited SHP2 in a quadruple unit cell of the 4DGP structure. (F) PMF of the binding cleft opening, d_cleft_, for autoinhibited SHP2 in the crystal environment.

Notably, the PMF further shows that the binding cleft is highly responsive to external perturbations, as only a small energy of 6–10 kJ/mol is required to close the binding cleft into a configuration found in crystal structures of autoinhibited SHP2 with well-resolved loops (e.g 4DGP.A, 4OHH.A, 5I6V.B, cf. Fig. 3) [16, 49, 50]. Therefore, the EF and the BG loop lining the N-SH2 binding cleft are constitutively flexible, and the binding cleft rearrangement in isolated N-SH2 requires only a small amount of free energy.

### Binding cleft opening is favored in autoinhibited SHP2 in water

In order to reveal the effect of the PTP allosteric interaction on the N-SH2 binding cleft, we computed the PMF of the binding cleft opening for SHP2 in water, again along the distance *d*_cleft_ (Figure 5C/D). Critically, for all umbrella windows, the C_*α*_–C_*α*_ distance *d*_block_ between Asp^61^ (belonging to N-SH2 blocking loop) and Ala^460^ (belonging to PTP catalytic loop) fluctuated around 6 Å (Figure S3), mainly below the experimental distances observed in autoinhibited structures of SHP2 (Figure 4B). Therefore, the N-SH2 domain remained closely bound to PTP throughout calculations, and the final PMF in Figure 5D represents the opening of the binding cleft in autoinhibited SHP2.

Surprisingly, even in tight interaction with PTP, a wide-open binding cleft of N-SH2 corresponds to the free energy minimum (Figure 5D). This result disagrees with the “allosteric switch” model [32, 39], which instead expects a closed and inaccessible binding cleft when the N-SH2 is bound to PTP. In addition, the PMF shows that a relatively small amount of free energy of 3–9.5 kJ/mol is required to close the binding cleft towards conformations found in crystal structures (Fig. 5D, black dots). This suggests that, on one hand, the binding cleft remains flexible and susceptible to structural rearrangements in case of ligand binding. On the other hand, the energy involved in binding cleft rearrangement is presumably many times smaller than the energy required to destabilize the N-SH2/PTP interface, promoting SHP2 activation. In fact, previous molecular dynamics simulations suggested that the complete displacement of N-SH2 away from the PTP active site requires an energy in the order of 70 kJ/mol [41, 43]. Hence, it is unlikely that rearrangement of the binding cleft are sufficient to drive SHP2 activation.

Taken together, the PMF shows that (i) even in complex with PTP, the binding cleft of N-SH2 is mostly open and accessible to ligands, and (ii) the binding cleft is not tensed and, therefore, rearrangement of the binding cleft alone are insufficient to drive SHP2 activation.

### Crystal contacts hinder the binding cleft opening

The PMF of SHP2 in solution suggests that the binding cleft is more open in solution than expected from available crystal structures (previous paragraph). In order to shed light on the origin of this discrepancies, we computed the PMF of binding cleft opening for SHP2 in the crystal environment of the 4DGP structure, for which the loops have been well resolved (Figure 5E, cf. Fig. 3). Simulations of a protein in the crystal reveals the influence of the crystal packing on protein structure and dynamics by comparing the results with simulations performed with the same system in solution. Such comparisons are still rare in the literature because researchers are often more interested in the properties of a protein in aqueous solution, corresponding to physiological conditions or most experiments [51–54].

The PMF reveals a clear effect of the crystal environment on the binding cleft conformation (Figures 5D and 5F). Namely, the most stable conformation in the crystal corresponds more closely to the crystal structure with a closed binding cleft (i.e. 4DGP) [16], whereas the binding cleft opening is strongly hindered. Hence, the tendency of the binding cleft to adopt a closed and inaccessible conformation in crystal structures is likely a bias due to the crystal contacts between protein replicas, and not a consequence of an allosteric mechanism imposed by N-SH2 binding to the PTP active site.

### Binding cleft opening does not promote the activation of SHP2

Based on the observation that the closed binding cleft is found in crystal structures in correspondence of the autoinhibited SHP2, whereas the crystal structures of the isolated, peptide-bound N-SH2 exhibited a full open, accessible binding cleft, it was hypothesized that the EF loop, together with the BG loop, may serve as allosteric switch in SHP2 activation [32, 39]. According to the “allosteric switch” model, the interaction with the PTP active site forces the N-SH2 binding cleft into a closed, inaccessible conformation, that prevents ligand binding (switch off); binding of a phosphopeptide would push the N-SH2 binding cleft towards an open, fully accessible conformation, with the consequence of weakening the interactions between the blocking loop and the catalytic site, promoting the N-SH2 domain release from PTP (switch on) [32, 39].

To definitely test such a model, we used umbrella sampling to compute the PMF of SHP2 activation (Figure 6A) with the N-SH2 binding cleft restrained either in closed (switch off) or in open (switch on) conformation. As reaction coordinate for SHP2 activation, we used the center-of-mass distance between the backbone atoms of the blocking loop and the catalytic PTP loop. Upon pulling the simulation system along this coordinate, the N-SH2 domain moved from its position in the autoinhibited state to different positions on the PTP surface, typically by sliding over the PTP surface. Among independent simulation runs, the final position of N-SH2 differed, suggesting that activated SHP2 covers a large conformational space.

**Figure 6:**
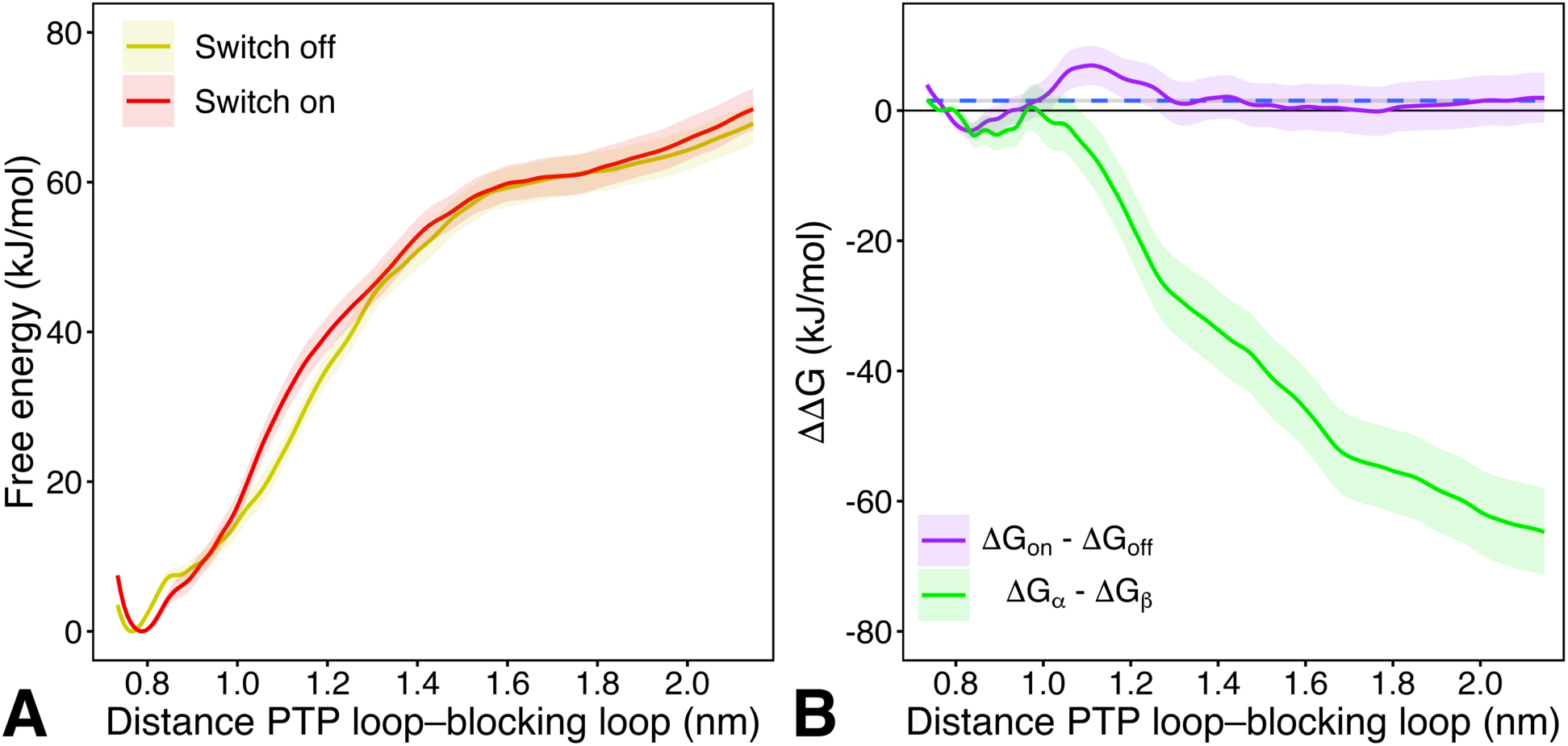
(A) PMFs of SHP2 opening with the N-SH2 binding cleft restrained either in closed (switch off, yellow line) or in open (switch on, red line) conformation. The distance between the blocking loop and the catalytic PTP loop, in terms of distance between the backbone centers-of-mass of residues 60-62 and residues 460-462, was taken as reaction coordinate. (B) ΔΔG profiles of SHP2 opening, calculated either for the “switch on” conformation respect to the “switch off” conformation (violet line), or for the α-state respect to the β-state (green line). Dashed blue line represents the average ΔΔG value of the ΔG_on_ – ΔG_off_ profile.

In disagreement with the “allosteric switch” model, the two PMF for “switch off” (Figure 6A, yellow line) and “switch on” (Figure 6A, red line), are almost identical, demonstrating that a switch of the binding cleft does not drive dissociation of the N-SH2 domain from PTP. The only perceptible, small difference at proximal distances is the slight right-shift of the “switch on” PMF minimum, likely ascribable to the different tilt of the helix *α*B, whose effects have been predicted from crystal structure analysis [32]. However, such perturbations do not further compromise the interface of the complex at larger distances.

In Figure 6B we reported the ΔΔG difference between the two profiles (ΔG_on_ – ΔG_off_). The flat profile (Figure 6B, violet line), whose average is virtually null (blue dashed line), confirms that the binding cleft conformation alone has virtually no influence on SHP2 activation. To test if instead the *α*–*β* model rationalizes SHP2 activation, ΔΔG profile of activation was computed with the N-SH2 domain either restrained in the *α*-or in the *β*-state (Figure 6B, green line). Evidently, the ΔΔG values become more and more negative as the distance between N-SH2 and PTP increases, demonstrating that the release of N-SH2 is greatly favored by the *α*-state.

In conclusion, PMFs indicated that the propensity of the N-SH2 domain to release PTP is essentially unresponsive to the conformation adopted by its binding cleft. Instead, the opening of the central *β*-sheet, as adopted by the *α*-state, leads to a larger rearrangement of the N-SH2 shape, therefore promoting SHP2 activation.

## DISCUSSION AND CONCLUSIONS

The conformational transition of disordered regions, occurring upon ligand binding or chemical modifications, is a recurrent molecular mechanism in protein regulation [42,55,56], which is why, for a long time, the flexible loops in the binding cleft of N-SH2 have been considered as molecular switches in an allosteric mechanism controlling SHP2 activation [39, 42]. However, in this study we have brought several arguments in opposition to the “allosteric switch” model: *i*) the structural disorder of the N-SH2 loops in crystal structures, *ii*) the constitutive flexibility of the binding cleft in solution, *iii*) the unexpected opening of the binding cleft, occurring even in autoinhibited SHP2 in water, *iv*) the steric hindrance, induced by the crystal contacts, forcing the binding cleft into a closed conformation, and finally *v*) the missing destabilization of the N-SH2/PTP interface upon binding cleft opening.

Experimental evidence brought up in favor of the “allosteric switch” model stated that *i*) the phosphopeptide binding is generally stronger to the isolated N-SH2 domain than to the N-SH2 domain as part of SHP2 [21], *ii*) the mutations destabilizing the N-SH2/PTP interface not only increase the SHP2 basal activity, but also result in a larger phosphopeptide binding affinity, albeit these mutations do not cluster at the N-SH2 binding site and apparently they do not affect the structure of N-SH2 [40]. Thus, the direct correlation between the basal activity of SHP2 and the ligand binding affinity to N-SH2, seems coherent with the “allosteric switch” model, because an eventual enhancement of the binding cleft opening would consequently promote the ligand binding to N-SH2. However, such experimental evidence would be potentially in agreement with any other allosteric mechanism involving an equilibrium between a stabilizing conformation and any activating conformation of N-SH2, independently on the structural details governing such an equilibrium. As a matter of fact, the assumption of an equilibrium between a stabilizing conformation, blocked by the interaction with PTP, and an activating conformation, induced by the binding of a phosphopeptide, inevitably leads to the conclusion that any weakening of the N-SH2/PTP interaction not only penalizes the stabilizing conformation in autoinhibited SHP2, but also indirectly makes N-SH2 more prone to bind a ligand. Such a conclusion is valid in case of the “allosteric switch” model, as well as for the *α*–*β* model.

Certain gain-of-function pathogenic mutations in SHP2 do not cluster at the N-SH2/PTP interface, where they could destabilize the N-SH2/PTP interactions, but seem to have more subtle, allosteric effects, since their impact on SHP2 function cannot be rationalized by mere steric effects. A typical example is the Noonan Syndrome (NS)-causing Thr42Ala substitution that replaces a conserved threonine in the central *β*-sheet [15]. Because Thr^42^ forms an H-bond with the phosphotyrosine in wild-type N-SH2, one might expect that Thr^42^ contributes to the stability of the N-SH2/phosphopeptide complex. However, the substitution with alanine leads to an increase in phosphopeptide binding affinity [15], as documented by the dramatically enhanced catalytic activity of the SHP2^A42^ mutant upon stimulation with a bisphosphoryl tyrosine-based activation motif (BTAM) peptide [15]. Notably, the “allosteric switch” model has been so far unable to convincingly explain the pathogenicity of the Thr42Ala substitution, in terms of either enhancement of the binding cleft opening or formation of additional favorable contacts with the phosphopeptide [15]. Instead, owing to the reduced propensity of alanine for *β*-sheet as compared to threonine [57], the *α*–*β* model attributes to the Thr42Ala substitution the effect of destabilizing the central *β*-sheet, thereby rendering N-SH2 more prone to adopt the activating *α*-state [43].

Analogously to the opening/closure of the binding cleft, the opening and closure of the central *β*-sheet in N-SH2 is detectable by comparing the crystal structures of autoinhibited SHP2 with crystal structures of the isolated N-SH2 domain bound to a phosphopeptide [31,32]. Thus, it does not come as a surprise that the opening of the central *β*-sheet results for the *α*–*β* model as the key determinant of the N-SH2 reshaping, which leads to the loss of complementarity for PTP, and thereby to the activation of SHP2. However, according to the *α*–*β* model, a coupling exists between the opening of the central *β*-sheet and the closure of either the pY loop or of the binding cleft at the +5 site. Therefore, we can conclude that the role of the binding cleft in N-SH2 is not to directly cause the loss of complementarity for PTP, but to induce, through the recognition and clamping of an activating phosphopeptide, a more extensive, concerted rearrangement in N-SH2 structure, which ends with the central *β*-sheet opening.

In conclusion, the *α*–*β* model includes, among others, also structural elements that were previously considered by the “allosteric switch” model. As such, the *α*–*β* model can be considered as a further generalized formulation of an allosteric mechanism, leading to a better comprehension of the structural transitions governing the activation of SHP2.

## METHODS

### Analyses on crystallographic structures

Structures of autoinhibited SHP2, in its wild-type form or mutants, were taken from RCSB Protein Data Bank: 2SHP [32], 4DGP [16], 4DGX [16], 4GWF, 4H1O, 4H34, 4NWF, 4NWG, 4OHD [50], 4OHE [50], 4OHH [50], 4OHI [50], 4OHL [50], 5I6V [49], 5IBM [49], 5IBS [49]. Dihedral angles were validated using the MolProbity web server [44]. Electron densities were inspected with Coot [58]. R-values and Z-scores were obtained from the residue-wise electron density using Uppsala Electron Density Server [48].

### MD simulations of the N-SH2 domain in solution

The initial atomic coordinates (residues 3-103) were derived from crystallographic structure 1AYD [31]. N-SH2 was put at the center of a dodecahedron box, large enough to contain the domain and at least 0.9 nm of solvent on all sides, and then it was solvated with 5577 explicit TIP3P [59] water molecules.

MD simulations were performed with the GROMACS software package [60], using the AMBER99SB*-ILDNP force field [61, 62] if not stated otherwise. Long-range electrostatic interactions were calculated with the particle-mesh Ewald (PME) approach [63]. A cutoff of 1.2 nm was applied to the direct-space Coulomb and Lennard-Jones interactions. Bond lengths and angles of water molecules were constrained with the SETTLE algorithm [64], and all other bonds were constrained with LINCS [65]. The pressure was set to 1 bar using the Parrinello-Rahman barostat [66]. The temperature was controlled at 300K using velocity-rescaling with a stochastic term [67].

The solvent was relaxed by energy minimization followed by 100 ps of MD at 300 K, while restraining protein coordinates with a harmonic potential. The system was then minimized without restraints and its temperature thermalized to 300 K in 10 ns, in a stepwise manner. After an equilibration run of 100 ns while N-SH2 was left completely unrestraint, the protein was equilibrated keeping the binding cleft closed. The C*_α_* distance between Gly^67^ (belonging to the EG loop) and Lys^89^ (belonging to the BG loop) was chosen as reaction coordinate, denoted *d*_cleft_-An umbrella potential (k = 10000 kJ/mol nm^2^) was progressively introduced in 10 ns, and maintained for other 100 ns, so that the *d*_cleft_ was finally constrained at 5 Å.

To maintain the N-SH2 domain with the central *β*-sheet parallel and closed, a flat-bottom potential (k = 10000 kJ/mol nm^2^), was introduced between the carbonyl C atom of Gly^39^ and the backbone N atom of Asn^58^, whose equilibrium distance was 4 Å; such a flat-bottom potential, acting only for distances larger than 4.4 Å, remained inactive during most part of the simulations, and it simply prevented the N-SH2 domain, rarely reaching the transition state, from further relaxing in a conformation with the central *β*-sheet completely open.

The reference distance of *d*_cleft_ = 7 Å was considered for the closed binding cleft (corresponding to the distance in 2SHP [32] crystal structure of autoinhibited SHP2), while a reference distance of 12 Å was chosen for the open binding cleft (corresponding to the distance in 1AYA [31] crystal structure of N-SH2 complexed with a phosphopeptide). Pulling simulation [68, 69] was carried out for gradually opening the binding cleft over 180 ns. The pull force constant was set to 1000 kJ mol^-1^nm^-2^, whereas the reference *d*_cleft_ was gradually increased from 5 to 14 Å with a velocity of 5×10^-6^ nm/ps.

To further improve the conformation sampling during these pulling simulations and to obtain more independent starting frames for umbrella sampling (see below), we coupled the pulling simulations with simulated tempering [70]. In line with previous studies [70, 71], simulated tempering simulations were carried out at constant volume, and temperature steps of 10 K between 300 and 380 K were applied. Temperature changes were controlled according to the Metropolis algorithm [72], to obtain canonical ensembles at all temperatures. The initial weights were chosen following Park and Pande [71]. Temperature transitions were attempted every 1 ps, and the weights were updated throughout the whole simulation according to the Wang-Landau adaptive weighting scheme [73].

The PMFs were obtained by means of umbrella sampling and WHAM [74, 75], using 46 windows, equispaced by 0.2 Å. Initial configurations for umbrella sampling were taken from the previous simulated tempering pulling simulations by means of a cluster analysis, thereby ensuring that the umbrella simulations were triggered from the most representative conformations. Accordingly, the configurations were collected from the sub-ensemble at 300 K, divided in 46 groups on the base of the umbrella window interval they belong, and clustered using the GROMOS clustering method [76]. The clustering cutoff was chosen on the base of the root mean-square deviation distribution in each group, picking the value corresponding to the first relative maximum in abscissa. The umbrella force constant was set to 1000 kJ mol^-1^nm^-2^, and a sampling of 1.5 *μ*s was performed for each window. Statistical errors of the PMFs were estimated with the Bayesian bootstrap of complete histograms [74], thereby considering only complete histograms as independent, yielding a maximum error of less than 2 kJ/mol.

### MD simulations of SHP2 in solution

The initial coordinates of SHP2 were taken from the autoinhibited conformation (2SHP) [32]. The protein was positioned at the center of a dodecahedral box, large enough to contain the protein and at least 0.9 nm of solvent on all sides, and solvated with ~23,000 explicit water molecules [59] and three Na^+^ ions. The structure was minimized and thermalized to 300 K using the procedure described above for N-SH2. A simulation of 1 *μ*s was performed to ensure a well-equilibrated autoinhibited structure in solution.

Next, the protein was equilibrated maintaining the binding cleft respectively either closed or open. An umbrella potential on *d*_cleft_ (k = 10000 kJ/mol nm^2^) was progressively introduced within 100 ns, and maintained for other 100 ns, so that *d*_cleft_ was finally constrained either at 5 Å or at 14 Å. The procedure was repeated four times providing same number of independent equilibrated configurations of SHP2 with the N-SH2 binding cleft either in closed or in open conformation. To avoid the opening of the central *β*-sheet in the N-SH2 domain, a flat-bottom potential was applied as described above.

Four pulling simulations were spawned either for gradually opening or for closing the binding cleft over 180 ns (eight simulations in total). The pull force constant was set to 1000 kJ mol^-1^nm^-2^, whereas the reference *d*_cleft_ was changed with a velocity of 5×10^-6^ nm/ps. Simulated tempering [70] was used to enhance sampling, using a temperature range from 300 to 380 K in steps of 5 K. In order to avoid the premature activation of SHP2 at higher temperatures and to maintain its autoinhibited structure, a flat-bottom potential (k = 10000 kJ/mol nm^2^), was introduced between the C*_α_* atom of Asp^61^ (belonging to the N-SH2 blocking loop) and the C*_α_* atom of Ala^460^ (belonging to PTP catalytic loop); this flat-bottom potential, only acting for distance *d*_block_ larger than 6.4 Å, was removed after the end of the simulated tempering pulling simulations. The PMF along *d*_cleft_ was computed as described above, except that each umbrella window was simulated for 250 ns.

### MD simulations of SHP2 in crystal environment

The initial coordinates of SHP2 were taken from the autoinhibited conformation (4DGP) [16]. This SHP2 crystal, which belongs to the P2_1_2_1_2 space group containing four symmetry-related molecules, has the following orthorhombic unit cell parameters: a = 5.485 nm, b = 22.110 nm, c = 4.036 nm, and *α* = *β* = *γ* = 90°. Missing or incomplete residues (strands Leu^236^–Gln^245^, Gly^295^–Val^301^, Phe^314^–Pro^323^, residues Met^1^, Thr^2^, Lys^235^, Arg^527^, Arg^528^ and the His-tagged tail Leu^529^–His^536^) were modelled by MOE [77] using the AMBER12:EHT force field and enabling the periodic system for P2_1_2_1_2 space group.

The starting coordinates of the single crystal unit cell were obtained by applying the P2_1_2_1_2 symmetry transformation. Next, the quadruple unit cell was generated by shifting the entire box along the x axis and/or z axis, resulting in an orthorhombic simulation box, with edges of 10.970, 22.110, and 8.072 nm, containing sixteen protein molecules. The use of a multiple unit cell as simulation box had the aim to forbid spurious couplings between the motion of the protein and its periodic images and to reduce unrealistic long-range correlations.

The AMBER99SB*-ILDNP force field [61, 62] was used, while a flat-bottom potential (k = 10000 kJ/mol nm^2^), was introduced between the carbonyl C atom of Gly^39^ and the backbone N atom of Asn^58^. The system was solvated with 26908 explicit water molecules [59] and 32 Na^+^ ions. The solvent box was generated by two successive solvent additions, each followed by a solvent relaxation session. The solvent was relaxed by energy minimization followed by 100 ps MD at 300 K, while restraining protein positions with a harmonic potential. The system was then energy-minimized without restraints and its temperature brought in a stepwise manner to 300 K in 10 ns.

In order to remove the conformational correlation between protein replicas, an equilibration run of 50 ns at 300 K was performed, followed by 50 ns of simulated tempering [70], with temperature ranging from 300 to 360 K in steps of 2 K, and another equilibration run of 50 ns.

Starting from the last system structure, two pulling simulations were spawned for either gradually opening or for gradually closing the binding cleft in each of the 16 protein replicas independently by others (32 simulations in total). The pull force constant was set to 1000 kJ mol^-1^nm^-2^, whereas the reference *d*_cleft_ was changed with a velocity of 5×10^-6^ nm/ps. Simulated tempering [70] was used to enhance sampling, using a temperature range from 300 to 360 K in steps of 2 K. Again, a flat-bottomed potential was used to avoid the opening of the central *β*-sheet, and immediately removed after the end of the simulated tempering pulling simulations. The PMF along *d*_cleft_ was computed as described above, except that each umbrella window was simulated for 20 ns.

### MD simulations of the activation of SHP2, with the binding cleft restrained into the closed or open conformation

The initial coordinates of SHP2 were taken from the autoinhibited conformation (2SHP) [32]. The protein was positioned at the center of a dodecahedral box, and solvated with ~23,000 explicit water molecules [59] and three Na^+^ ions. Because certain Amber force fields in conjunction with TIP3P have been reported to overstabilize protein-protein contacts [78], we used the CHARMM36m force field [79] for simulations of SHP2 activation. Hydrogen atoms were described as virtual sites, allowing a time step of 4 fs [80]. The structure was minimized and thermalized to 300 K using the procedure described above. A simulation of 1 *μ*s was performed to ensure a well-equilibrated autoinhibited structure in solution.

Next, the binding cleft of the N-SH2 domain was restrained either in closed (*d*_cleft_ = 5 Å) or in open conformation (*d*_cleft_ = 12 Å), using a harmonic restraint along *d*_cleft_ (k = 1000 kJ/mol nm^2^). Opening of the central *β*-sheet was excluded using a flat-bottomed potential, as described above. N-SH2 was pulled away from PTP using 600 ns pulling simulations at constant pull velocity [68]. The center-of-mass distance between the blocking loop (residues 60-62) and the catalytic PTP loop (residues 460-462) was chosen as reaction coordinate. That distance was pulled from 0.73 nm to 2.15 nm. To obtain statistically independent pathways for SHP2 opening, eight pulling simulations were carried out for each N-SH2 binding cleft state, restrained either in closed or in open conformation (16 simulations in total). To further improve the conformation sampling, we coupled the pulling simulations with simulated tempering [70], using a temperature range from 300 to 400 K in steps of 5 K.

The PMFs for the opening of SHP2 were obtained using umbrella sampling and the weighted histogram analysis method (WHAM) [74, 75]. Two PMFs were computed with the N-SH2 binding cleft either restrained in closed or in open conformation. For each profile, 72 windows, equispaced by 0.2 Å, were used. Initial configurations for umbrella sampling were taken from the previous simulated tempering pulling simulations by means of a cluster analysis [76]. The umbrella force constant was set to 4000 kJ mol^-1^nm^-2^, and sampling of 100 ns was performed for each window. Statistical errors of the PMFs were estimated using the Bayesian bootstrap of complete histograms [74].

## Data availability

All data regarding the work presented here is available upon request to the corresponding author.

## Acknowledgements

This study has received funding from the European Union’s Horizon 2020 research and innovation programme under the Marie Skłodowska-Curie grant agreement MARS n. 705829. This study was supported by the Deutsche Forschungsgemeinschaft (J.S.H., grant no 1971-4/1).

## Author contributions

All authors contributed to the final manuscript.

## References

[1] Tajan, M., de Rocca Serra, A., Valet, P., Edouard, T. & Yart. SHP2 sails from physiology to pathology. European Journal of Medical Genetics 58, 509–525 (2015).

[2] Dance, M., Montagner, A., Salles, J. P., Yart, A. & Raynal, P. The molecular functions of Shp2 in the Ras/Mitogen-activated protein kinase (ERK1/2) pathway. Cell Signal 20, 453–9 (2008).

[3] Shen, D. et al. Therapeutic potential of targeting SHP2 in human developmental disorders and cancers. Eur J Med Chem 190, 112117 (2020).

[4] Tang, K., Jia, Y. N., Yu, B. & Liu, H. M. Medicinal chemistry strategies for the development of protein tyrosine phosphatase SHP2 inhibitors and PROTAC degraders. Eur J Med Chem 204, 112657 (2020).

[5] Li, S. M. The biological function of shp2 in human disease. Mol Biol (Mosk) 50, 27–33 (2016).

[6] Digilio, M. C. et al. Grouping of multiple-lentigines/LEOPARD and Noonan syndromes on the PTPN11 gene. Am J Hum Genet 71, 389–94 (2002).

[7] Tartaglia, M. et al. Diversity and functional consequences of germline and somatic PTPN11 mutations in human disease. Am J Hum Genet 78, 279–90 (2006).

[8] Tartaglia, M. et al. Mutations in PTPN11, encoding the protein tyrosine phosphatase SHP-2, cause Noonan syndrome. Nat Genet 29, 465–8 (2001).

[9] Keilhack, H., David, F. S., McGregor, M., Cantley, L. C. & Neel, B. G. Diverse biochemical properties of Shp2 mutants. Implications for disease phenotypes. J Biol Chem 280, 30984–93 (2005).

[10] Qiu, W. et al. Structural insights into Noonan/LEOPARD syndrome-related mutants of protein-tyrosine phosphatase SHP2 (PTPN11). BMC Struct Biol 14, 10 (2014).

[11] Martinelli, S. et al. Counteracting effects operating on Src homology 2 domain-containing protein-tyrosine phosphatase 2 (SHP2) function drive selection of the recurrent Y62D and Y63C substitutions in Noonan syndrome. J Biol Chem 287, 27066–77 (2012).

[12] Edouard, T. et al. Functional effects of PTPN11 (SHP2) mutations causing LEOPARD syndrome on epidermal growth factor-induced phosphoinositide 3-kinase/AKT/glycogen synthase kinase 3beta signaling. Mol Cell Biol 30, 2498–507 (2010).

[13] Hanna, N. et al. Reduced phosphatase activity of SHP-2 in LEOPARD syndrome: consequences for PI3K binding on Gab1. FEBS Lett 580, 2477–82 (2006).

[14] Kontaridis, M. I., Swanson, K. D., David, F. S., Barford, D. & Neel, B. G. PTPN11 (Shp2) mutations in LEOPARD syndrome have dominant negative, not activating, effects. J Biol Chem 281, 6785–92 (2006).

[15] Martinelli, S. et al. Diverse driving forces underlie the invariant occurrence of the T42A, E139D, I282V and T468M SHP2 amino acid substitutions causing Noonan and LEOPARD syndromes. Hum Mol Genet 17, 2018–29 (2008).

[16] Yu, Z. H. et al. Structural and mechanistic insights into LEOPARD syndrome-associated SHP2 mutations. J Biol Chem 288, 10472–82 (2013).

[17] Ahmed, T. A. et al. SHP2 Drives Adaptive Resistance to ERK Signaling Inhibition in Molecularly Defined Subsets of ERK-Dependent Tumors. Cell Reports 26, 65–78.e5 (2019).

[18] Butterworth, S., Overduin, M. & Barr, A. J. Targeting protein tyrosine phosphatase SHP2 for therapeutic intervention. Future Medicinal Chemistry 6, 1423–1437 (2014).

[19] Chen, Y. N. et al. Allosteric inhibition of SHP2 phosphatase inhibits cancers driven by receptor tyrosine kinases. Nature 535, 148–52 (2016).

[20] Frankson, R. et al. Therapeutic targeting of oncogenic tyrosine phosphatases. Cancer Research 77, 5701 (2017).

[21] Hayashi, T. et al. Differential Mechanisms for SHP2 Binding and Activation Are Exploited by Geographically Distinct Helicobacter pylori CagA Oncoproteins. Cell Rep 20, 2876–2890 (2017).

[22] Hui, E. et al. T cell costimulatory receptor CD28 is a primary target for PD-1-mediated inhibition. Science 355, 1428–1433 (2017).

[23] Prahallad, A. et al. PTPN11 Is a Central Node in Intrinsic and Acquired Resistance to Targeted Cancer Drugs. Cell Re 12, 1978–85 (2015).

[24] Ran, H., Tsutsumi, R., Araki, T. & Neel, B. G. Sticking It to Cancer with Molecular Glue for SHP2. Cancer Cell 30, 194–196 (2016).

[25] Torres-Ayuso, P. & Brognard, J. Shipping Out MEK Inhibitor Resistance with SHP2 Inhibitors. Cancer Discov 8, 1210–1212 (2018).

[26] Tartaglia, M. et al. Somatic PTPN11 mutations in childhood acute myeloid leukaemia. Br J Haematol 129, 333–9 (2005).

[27] Tartaglia, M. et al. Genetic evidence for lineage-related and differentiation stage-related contribution of somatic PTPN11 mutations to leukemogenesis in childhood acute leukemia. Blood 104, 307–13 (2004).

[28] Goemans, B. F. et al. Differences in the prevalence of PTPN11 mutations in FAB M5 paediatric acute myeloid leukaemia. Br J Haematol 130, 801–3 (2005).

[29] Loh, M. L. et al. Acquired PTPN11 mutations occur rarely in adult patients with myelodysplastic syndromes and chronic myelomonocytic leukemia. Leuk Res 29, 459–62 (2005).

[30] Pandey, R., Saxena, M. & Kapur, R. Role of SHP2 in hematopoiesis and leukemogenesis. Curr Opin Hematol 24, 307–313 (2017).

[31] Lee, C. H. et al. Crystal structures of peptide complexes of the amino-terminal SH2 domain of the Syp tyrosine phosphatase. Structure 2, 423–38 (1994).

[32] Hof, P., Pluskey, S., Dhe-Paganon, S., Eck, M. J. & Shoelson, S. E. Crystal structure of the tyrosine phosphatase SHP-2. Cell 92, 441–50 (1998).

[33] LaRochelle, J. R. et al. Structural reorganization of SHP2 by oncogenic mutations and implications for oncoprotein resistance to allosteric inhibition. Nat Commun 9, 4508 (2018).

[34] Neel, B. G., Gu, H. & Pao, L. The ‘Shp’ing news: SH2 domain-containing tyrosine phosphatases in cell signaling. Trends Biochem Sci 28, 284–93 (2003).

[35] Liu, B. A. & Machida, K. Introduction: History of SH2 Domains and Their Applications. Methods Mol Biol 1555, 3–35 (2017).

[36] Kaneko, T., Joshi, R., Feller, S. M. & Li, S. S. Phosphotyrosine recognition domains: the typical, the atypical and the versatile. Cell Commun Signal 10, 32 (2012).

[37] Piccione, E. et al. Phosphatidylinositol 3-kinase p85 SH2 domain specificity defined by direct phosphopeptide/SH2 domain binding. Biochemistry 32, 3197–202 (1993).

[38] Marasco, M. & Carlomagno, T. Specificity and regulation of phosphotyrosine signaling through SH2 domains. J Struct Biol X 4, 100026 (2020).

[39] Barford, D. & Neel, B. G. Revealing mechanisms for SH2 domain mediated regulation of the protein tyrosine phosphatase SHP-2. Structure 6, 249–254 (1998).

[40] Bocchinfuso, G. et al. Structural and functional effects of disease-causing amino acid substitutions affecting residues Ala72 and Glu76 of the protein tyrosine phosphatase SHP-2. Proteins 66, 963–74 (2007).

[41] Darian, E., Guvench, O., Yu, B., Qu, C. K. & MacKerell, J., A. D. Structural mechanism associated with domain opening in gain-of-function mutations in SHP2 phosphatase. Proteins 79, 1573–88 (2011).

[42] Lim, W. A. The modular logic of signaling proteins: building allosteric switches from simple binding domains. Curr Opin Struct Biol 12, 61–8 (2002). URL https://www.ncbi.nlm.nih.gov/pubmed/11839491.

[43] Anselmi, M. & Hub, J. S. An allosteric interaction controls the activation mechanism of shp2 tyrosine phosphatase. Scientific Reports 10, 18530 (2020).

[44] Williams, C. J. et al. MolProbity: More and better reference data for improved all-atom structure validation. Protein Sci 27, 293–315 (2018).

[45] Laskowski, R. A. Structural quality assurance. Structural Bioinformatics 273–303 (2003).

[46] Pražnikar, J., Tomić, M. & Turk, D. Validation and quality assessment of macromolecular structures using complex network analysis. Scientific reports 9, 1678–1678 (2019).

[47] Tickle, I. J. Statistical quality indicators for electron-density maps. Acta crystallographica. Section D, Biological crystallography 68, 454–467 (2012).

[48] Kleywegt, G. J. et al. The Uppsala Electron-Density Server. Acta Crystallographica Section D 60, 2240–2249 (2004).

[49] LaRochelle, J. R. et al. Structural and Functional Consequences of Three Cancer-Associated Mutations of the Oncogenic Phosphatase SHP2. Biochemistry 55, 2269–77 (2016).

[50] Yu, Z. H. et al. Molecular basis of gain-of-function LEOPARD syndrome-associated SHP2 mutations. Biochemistry 53, 4136–51 (2014).

[51] Anselmi, M., Brunori, M., Vallone, B. & Di Nola, A. Molecular dynamics simulation of the neuroglobin crystal: comparison with the simulation in solution. Biophys J 95, 4157–62 (2008).

[52] Cerutti, D. S., Le Trong, I., Stenkamp, R. E. & Lybrand, T. P. Dynamics of the streptavidin-biotin complex in solution and in its crystal lattice: distinct behavior revealed by molecular simulations. J Phys Chem B 113, 6971–85 (2009).

[53] Vorontsov, I. & Miyashita, O. Solution and crystal molecular dynamics simulation study of m4-cyanovirin-N mutants complexed with di-mannose. Biophys J 97, 2532–40 (2009).

[54] Zhang, Z. & Wriggers, W. Polymorphism of the epidermal growth factor receptor extracellular ligand binding domain: the dimer interface depends on domain stabilization. Biochemistry 50, 2144–56 (2011).

[55] Laskowski, R. A., Gerick, F. & Thornton, J. M. The structural basis of allosteric regulation in proteins. FEBS Lett 583, 1692–8 (2009).

[56] Tsai, C. J., Del Sol, A. & Nussinov, R. Protein allostery, signal transmission and dynamics: a classification scheme of allosteric mechanisms. Mol Biosyst 5, 207–16 (2009).

[57] Minor, D. L. & Kim, P. S. Measurement of the *β*-sheet-forming propensities of amino acids. Nature 367, 660–663 (1994).

[58] Emsley, P., Lohkamp, B., Scott, W. G. & Cowtan, K. Features and development of Coot. Acta Crystallogr D Biol Crystallogr 66, 486–501 (2010).

[59] Jorgensen, W. L., Chandrasekhar, J., Madura, J. D., Impey, R. W. & Klein, M. L. Comparison of simple potential functions for simulating liquid water. J. Chem. Phys. 79, 926–935 (1983).

[60] Van Der Spoel, D. et al. GROMACS: fast, flexible, and free. J Comput Chem 26, 1701–18 (2005).

[61] Aliev, A. E. et al. Motional timescale predictions by molecular dynamics simulations: case study using proline and hydroxyproline sidechain dynamics. Proteins 82, 195–215 (2014).

[62] Best, R. B. & Hummer, G. Optimized molecular dynamics force fields applied to the helix-coil transition of polypeptides. J Phys Chem B 113, 9004–15 (2009).

[63] Darden, T., York, D. & Pedersen, L. Particle Mesh Ewald: an Nlog(N) method for Ewald sum in large systems. J. Chem. Phys. 98, 10089–10092 (1993).

[64] Miyamoto, S. & Kollman, P. A. SETTLE: An analytical version of the SHAKE and RATTLE algorithms for rigid water models.. J. Comp. Chem. 13, 952–962 (1992).

[65] Hess, B., Bekker, H., Berendsen, H. J. C. & Fraaije, J. G. E. M. LINCS: a linear constraint solver for molecular simulations. Journal of Computational Chemistry 18, 1463–1472 (1997).

[66] Parrinello, M. & Rahman, A. Polymorphic transitions in single crystals: A new molecular dynamics method. J. Appl. Phys. 52, 7182–7190 (1981).

[67] Bussi, G., Donadio, D. & Parrinello, M. Canonical sampling through velocity rescaling. J. Chem. Phys. 126, 14101 (2007).

[68] Grubmuller, H., Heymann, B. & Tavan, P. Ligand binding: molecular mechanics calculation of the streptavidin-biotin rupture force. Science 271, 997–9 (1996).

[69] Izrailev, S. et al. Steered molecular dynamics, 39–65 (Springer-Verlag, 1998).

[70] Marinari, E. & Parisi, G. Simulated tempering: a new Monte Carlo scheme. Europhys. Lett. 19, 451–458 (1992).

[71] Park, S. & Pande, V. S. Choosing weights for simulated tempering. Phys Rev E Stat Nonlin Soft Matter Phys 76, 016703 (2007).

[72] Metropolis, N., Rosenbluth, A. W., Rosenbluth, M. N. & Teller, A. H. Equation of state calculations by fast computing machines. J. Chem. Phys. 21, 1087 (1953).

[73] Wang, F. & Landhow to insertau, D. P. Efficient, multiple-range random walk algorithm to calculate the density of states. Phys Rev Lett 86, 2050–3 (2001).

[74] Hub, J. S., de Groot, B. L. & van der Spoel, D. g_wham—A Free Weighted Histogram Analysis Implementation Including Robust Error and Autocorrelation Estimates. Journal of Chemical Theory and Computation 6, 3713–3720 (2010).

[75] Kumar, S., Bouzida, D., Swendsen, S. H., Kollman, P. A. & Rosenberg, J. M. The Weighted Histogram Analysis Method for free-energy calculations on biomolecules. I. the method. Journal of Computational Chemistry 13, 1011–1021 (1992).

[76] Daura, X. et al. Peptide Folding: When Simulation Meets Experiment. Angewandte Chemie International Edition 38, 236–240 (1999).

[77] Chemical Computing Group Inc. Molecular Operating Environment (MOE) software (2014).

[78] Best, R. B., Zheng, W. & Mittal, J. Balanced protein–water interactions improve properties of disordered proteins and non-specific protein association. Journal of Chemical Theory and Computation 10, 5113–5124 (2014).

[79] Huang, J. et al. CHARMM36m: an improved force field for folded and intrinsically disordered proteins. Nat Methods 14, 71–73 (2017).

[80] Feenstra, K. A., Hess, B. & Berendsen, H. J. C. Improving efficiency of large time-scale molecular dynamics simulations of hydrogen-rich systems. Journal of Computational Chemistry 20, 786–798 (1999).

