## Supporting information for "Do the loops in the N-SH2 binding cleft truly serve as allosteric switch in SHP2 activation?"

M.A. 0000-0001-5711-770X

J.S.H. 0000-0001-7716-1767

* Corresponding author:

 (MA)

KEYWORDS: Molecular dynamics simulations, Phosphatases, Protein dynamics, Protein function, Allosteric regulation

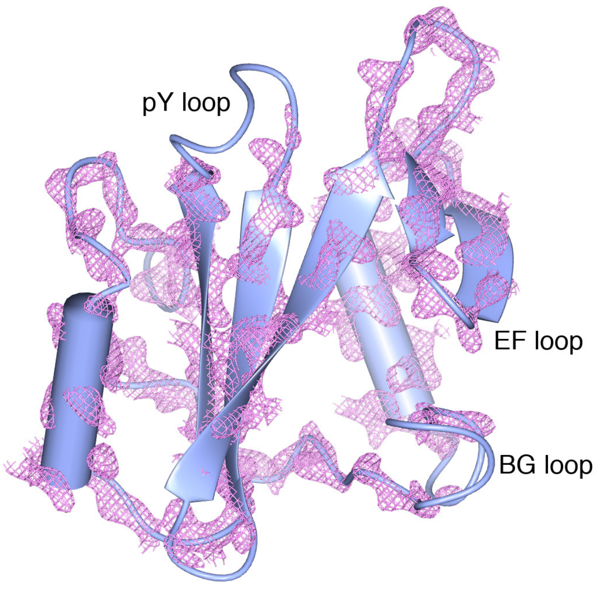


Figure S1: Electron density map of the N-SH2 domain backbone as obtained from the crystal structure of autoinhibited SHP2 (pdbID 2SHP) [1]. The positions of the flexible loops (pY loop, EF loop, and BG loop) are indicated by the labels.

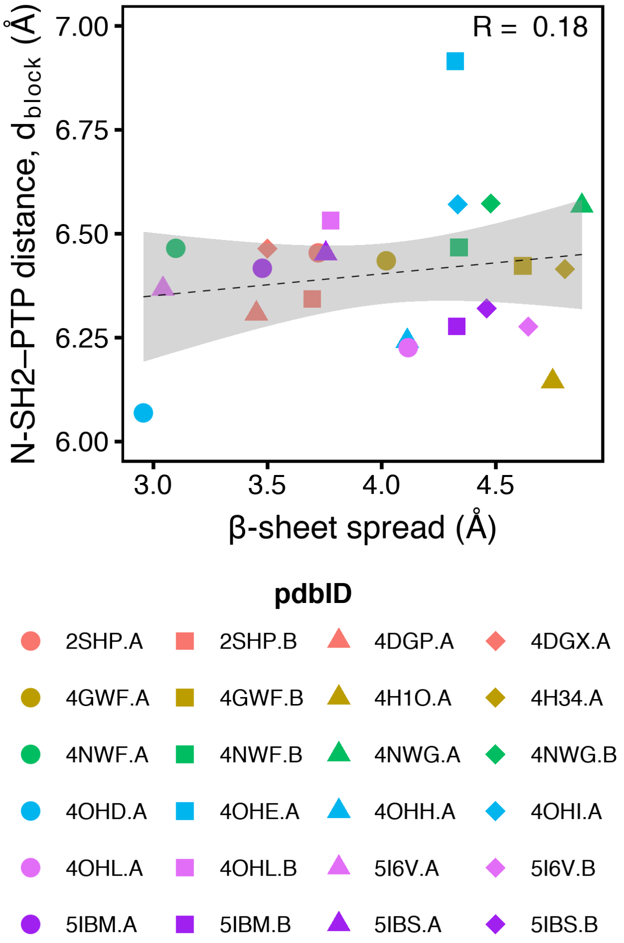


Figure S2: Correlation between the N-SH2 central $\beta$-sheet spread and the N-SH2 blocking loop distance from the catalytic PTP loop, $d_{\mathrm{block}}$ (Asp$^{61}$ C$_\alpha$–Ala$^{460}$ C$_\alpha$ distance), as taken from crystal structure of autoinhibited SHP2, comprising either wild type or functional mutants [1–4].

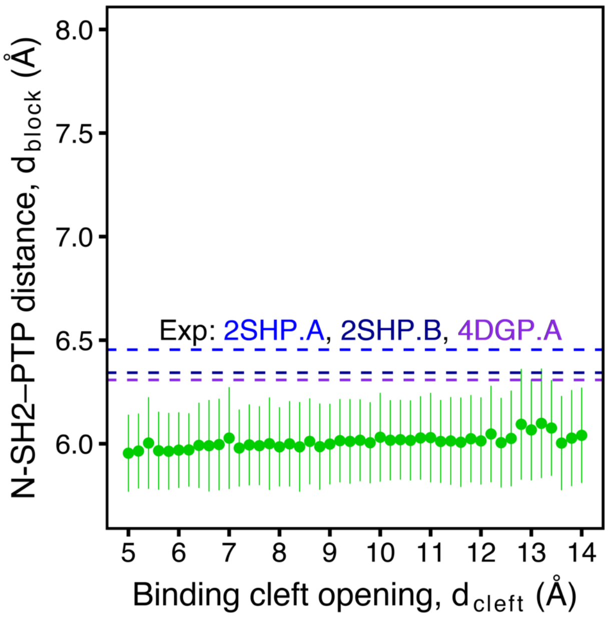


Figure S3: Dot plot of the N-SH2 blocking loop distance from the catalytic PTP loop, $d_{\text{block}}$ (Asp$^{61}$ $C_\alpha$–Ala$^{460}$ $C_\alpha$ distance), as a function of the N-SH2 binding cleft opening, $d_{\text{cleft}}$ (Gly$^{67}$ $C_\alpha$–Lys$^{89}$ $C_\alpha$ distance), as obtained from umbrella sampling simulations of autoinhibited SHP2 in water. The bars indicate the RMS fluctuation of $d_{\text{block}}$, that remains below the experimental $d_{\text{block}}$ distances observed in autoinhibited structures of wild-type SHP2 (pdbID 2SHP, 4DGP) [1, 2].

### References


[1] Hof, P., Pluskey, S., Dhe-Paganon, S., Eck, M. J. & Shoelson, S. E. Crystal structure of the tyrosine phosphatase SHP-2. *Cell* **92**, 441–50 (1998).

[2] Yu, Z. H. *et al.* Structural and mechanistic insights into LEOPARD syndrome-associated SHP2 mutations. *J Biol Chem* **288**, 10472–82 (2013).

[3] LaRochelle, J. R. *et al.* Structural and Functional Consequences of Three Cancer-Associated Mutations of the Oncogenic Phosphatase SHP2. *Biochemistry* **55**, 2269–77 (2016).

[4] Yu, Z. H. *et al.* Molecular basis of gain-of-function LEOPARD syndrome-associated SHP2 mutations. *Biochemistry* **53**, 4136–51 (2014).
